# Differential dopamine receptor-dependent sensitivity improves action selection in the basal ganglia

**DOI:** 10.1101/2020.11.12.380451

**Authors:** Olivier Codol, Paul L. Gribble, Kevin N. Gurney

**Affiliations:** Department of Psychology, University of Western Ontario, London, ON, Canada; The University of Sheffield, Sheffield, UK

**Author notes:** Correspondence to: Olivier Codol < >.

## Abstract

The problem of selecting one action from a set of different possible actions, simply referred to as the problem of action selection, is a ubiquitous challenge in the animal world. For vertebrates, the basal ganglia (BG) are widely thought to implement the core computation to solve this problem, as the anatomy and physiology of the BG are well-suited to this end. However, the BG still displays physiological features whose role in achieving efficient action selection remains unclear. In particular, it is known that the two types of dopaminergic receptors (D1 and D2) present in the BG give rise to mechanistically different responses. The overall effect will be a difference in sensitivity to dopamine which may have ramifications for action selection. However, which receptor type leads to a stronger response is, a priori, unclear, due to the complexity of the intracellular mechanisms involved. In this study, we use the action selection hypothesis to *predict* which of D1 or D2 has the greater sensitivity. Thus, we ask - what sensitivity ratio would result in enhanced action selection functionality in the basal ganglia? To do this, we incorporated differential D1 and D2 sensitivity in an existing, high level computational model of the macro-architecture of the basal ganglia, via a simple weighting variable. We then quantitatively assessed the model’s capacity to perform action selection as we parametrically manipulated the new feature. We show that differential (rather than equal) D1 and D2 sensitivity to dopaminergic input improves action selection, and specifically, that greater D1 sensitivity (compared to that for D2) leads to these improvements.

## 1. Action selection, the basal ganglia, and the role of dopamine

### 1.1. The basal ganglia as a central switch for action selection

The need for selection usually rises when several systems need to use a quantitatively limited resource. In the case of action selection, several candidate actions often require the same body effectors, making those actions mutually exclusive and requiring the brain to implement a solution to the ensuing action selection problem. Arguably the most efficient architecture to that end is that of a *central switch* (Redgrave et al., 1999), whereby competing actions are directly and reciprocally wired to a central unit implementing a competition mechanism. Indeed, a central switch allows for the number of required connections to increase at a much slower pace than alternative architectures as the number of potential actions increases. This is particularly important in the brain, where inter-region connectivity is strongly restricted by geometric considerations (Ringo, 1991). The notion that the basal ganglia (BG) act as the central switch in action selection is supported by both theoretical (Gurney et al., 2001a; Humphries et al., 2006) and physiological (Friend & Kravitz, 2014; Mink, 1996) evidence.

### 1.2. Overview of the basal ganglia architecture

The primate BG is composed of several distinct nuclei with a complex but reasonably well-understood connectivity. We use a somewhat simplified architecture which, nevertheless, captures much of this structure and connectivity (Mink, 1996). The principal nuclei are the striatum, the subthalamic nucleus (STN), the globus pallidus (GP), and the substantia nigra (SN; figure 1a). The GP is also sub-divided into the internal (GPi) and external (GPe) segments, and the SN into the *pars reticulata* (SNr) and *pars compacta* (SNc; figure 1b). The biggest proportion of the SNc neural population consists of dopaminergic (dopamine) neurons that send diffuse projections to the striatum (Smith et al., 1994). There, they connect to two different neuronal populations, each expressing a different subtype of dopamine receptors, called D1 and D2 receptors. Interestingly, D1-expressing neurons in the striatum project predominantly to the GPi/SNr while D2-expressing neurons project to the GPe (figure 1a), with all these striatal projections being inhibitory (Ericsson et al., 2013; Smith et al., 1998). Finally, unlike any other nucleus in the BG, the STN sends excitatory projections to its targets, which are the GPi/SNr and the GPe (Shink et al., 1996). Further, these projections are *diffuse* (Parent & Hazrati, 1993). In return, the GPe sends *focused,* rather than diffuse, reciprocal inhibitory projections back to the STN. It also sends focused inhibitory projections to the GPi/SNr.

**Figure 1.**
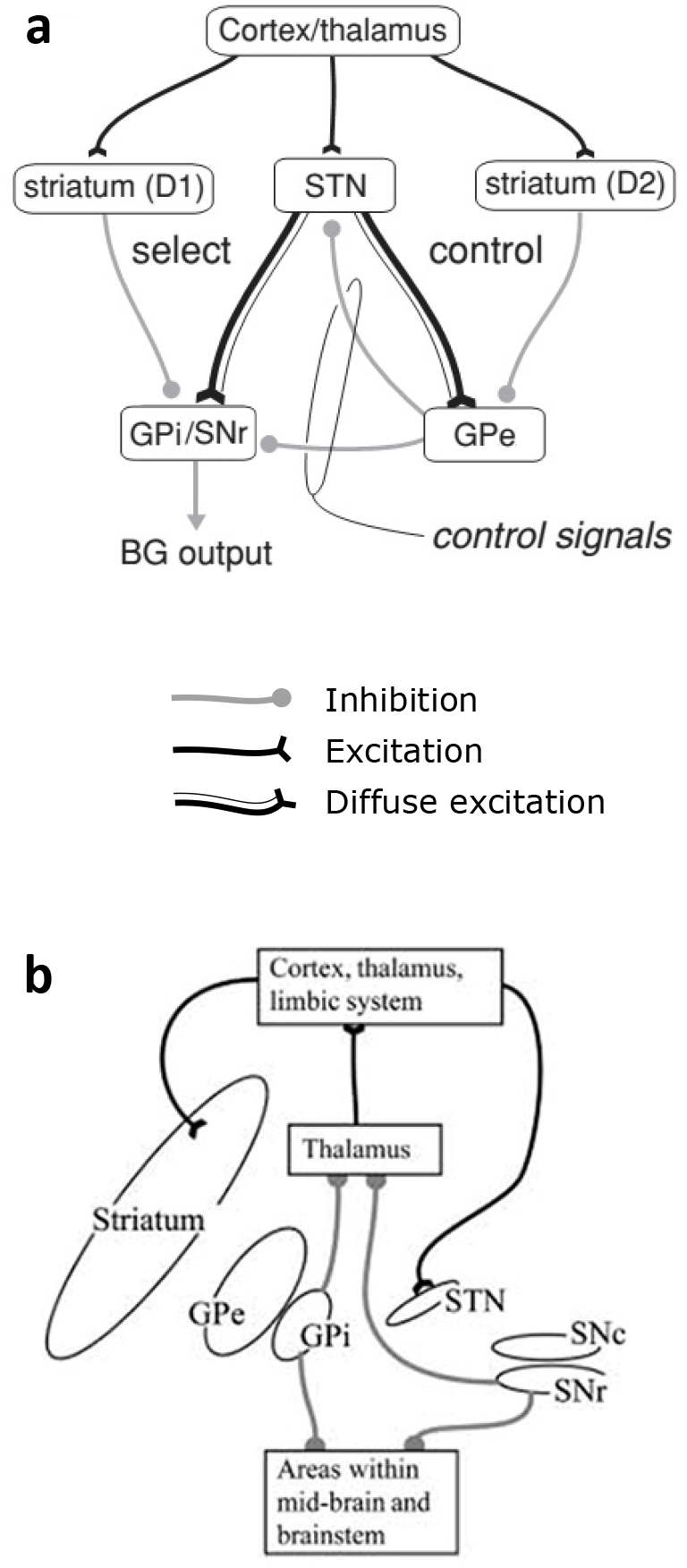
An overview of the anatomical architecture of the BG. **(a)** Intrinsic connectivity of the BG. **(b)** Extrinsic connectivity of the BG, showing the baso-thalamo-cortical loops. *Figure modified from Gurney et al. (2001a).*

Consistent with the action selection hypothesis and the central switch hypothesis, the BG receives a massive amount of projections from virtually the whole cortex (except the primary visual and auditory cortex), the thalamus and the limbic system (Coizet et al., 2009; Feger et al., 1994; Lanciego et al., 2004; Monakow et al., 1978; Nakano et al., 2000; Sharpe et al., 2019). Consequently, the BG can theoretically receive a great variety of sensory, cognitive, and motivational information (Sharpe et al., 2019), which provides the required contextual information to properly define the relative value of each action (*e.g.* prominence of the stimulus or immediate utility). These various inputs enter the BG through the striatum and the STN. The main output nuclei of the BG are the GPi and the SNr, which innervate the ventral thalamus, which in turn projects back to the cortex. This thalamo-cortical connection therefore completes an anatomical loop that goes from the cortex to the BG to the thalamus, and back to the cortex (Alexander et al., 1986; Wiesendanger et al., 2004). The BG contains a large amount of these loops, which are organized in parallel with each other. Critically, the BG’s default output is a tonic inhibition from the GPi/SNr to the thalamus (figure 1a, b). Actions are supposed to be permitted in BG by selective release of this inhibition so that reduced output from GPi/SNr leads to disinhibition of an action (Chevalier & Deniau, 1990).

### 1.3. The basal ganglia architecture supports the action selection hypothesis

The anatomy of the BG outlined here is well suited for action selection, or any selection computation in general (Bariselli et al., 2019; Kwak & Jung, 2019). This was illustrated by Gurney, Prescott and Redgrave in a computational model of the BG (Gurney et al., 2001a;0) — henceforth referred to as the “GPR” model — that reproduces the functional architecture in figure 1a. The main assumption of the GPR model is that each action is represented in the BG by a specific stream of sensory, cognitive and emotional information called a “channel” which represents one possible action to perform. Channels are organized in parallel, in line with physiological and histological evidence (Alexander et al., 1986; Graybiel et al., 1981). Those channels then compete with each other through off-center/on-surround computation in the BG circuit, with the “winning” channel eventually suppressing the other channels’ activity to promote its own selection (Girard et al., 2020).

Center/surround activity has been experimentally observed in the BG (Mink & Thach, 1993), and is the result of a diffuse excitatory drive from the STN and focused inhibitory drive from the striatum (figure 2a, b) (Gurney et al., 2001a). If a channel, say, the middle channel in figure 2b, receives a stronger drive from the cortex (figure 2b), the STN will more strongly activate all channels in the output layer through its diffuse projection, while only the middle channel will receive an equally strong inhibition from the striatum. Therefore, the middle channel’s output will be em less than the other channels’ output, as required by the disinhibition hypothesis outlined above.

As can be seen in figure 2b, a single channel in the STN will contact all channels in the output layer, which effectively leads to the on-surround competition illustrated in figure 2a. However, this also means, by extension, that each channel in the output layer receives excitatory input from all channels (figure 2c). Consequently, as the number of channels increases, the excitatory drive into each output channel will increase, while the inhibitory drive from the striatum will remain the same. Therefore, the total excitatory drive must be scaled to avoid runaway excitation as the number of channels increases. Ideally, to obtain this, each excitatory input from the STN should be weighted by 1/n, where *n* is the number of channels engaged. However, to achieve such a nuanced synaptic weight scaling is biologically challenging. An interesting alternative, which appears biologically attainable, is to provide an ‘automatic gain control’ to the STN to keep its overall drive in check (Gurney et al., 2001a;0). Interestingly, the BG appears to have a candidate mechanism for this in the circuit comprising GPe, STN and striatum (figure 1a). The GPe receives a similar combination of inputs to the GPi/SNr and then provides inhibition to the STN proportional to the drive supplied by the latter. We refer to this of normalization of activity as *capacity scaling* and it ascribes a new role to the GPe - that of supplying ‘control’ signals to the main selection network (comprising striatum, STN and GPi/SNr). It should be noted that this functional interpretation of the BG into a control/selection pathway diverges from the more classical view that considers a pathway dichotomy based on “direct” and “indirect” routes to the BG output instead (Albin et al., 1989; Bariselli et al., 2019; Kwak & Jung, 2019).

### 1.4. Dopamine and action selection

Dopaminergic input in the BG originates from the SNc, and exhibits both tonic and phasic activity patterns (Grace, 1991). However, phasic activity appears mainly to relate to reinforcement learning (Schultz, 1998) rather than action selection, so we will only consider tonic activity in this work. Tonic dopaminergic input is believed to modulate the action selection computation (Groves et al., 1994). This has been interpreted in the GPR/action selection context as allowing the BG to switch between two regimes (Gurney et al., 2004): one, which we call ‘hard selection’, promotes single-action selection, and another – ‘soft selection’ – allows multiple actions to be selected. The latter may be particularly relevant when competing actions are not strictly mutually exclusive (*e.g.* picking a fruit in each hand), or in learning situations where exploratory behaviour is required.

Striatal neurons contacted by dopaminergic projections are segregated into two distinct populations with different dopaminergic receptors, simply labelled D1 and D2. D1 receptors facilitate cortico-striatal transmission, whereas D2 receptors attenuate it (Surmeier et al., 2007). D1 and D2 receptors display quantitatively different molecular affinities to dopamine, with D2 receptors binding more easily to dopamine (90%) than D1 receptor (20%) (Arbuthnott & Wickens, 2007; Richfield et al., 1989). However, these numbers have been challenged by more recent studies and what, mechanistically, underlies binding affinities of D1 and D2 is still unresolved (Cumming, 2011; Skinbjerg et al., 2012). Additionally, the intracellular activation pathways differ between D1 and D2. Therefore, the overall response to dopamine at either receptor type will be the result of a complex cascade of influences, which may even work in opposing directions to the difference in binding affinity (Kenakin, 2013). It is not surprising, therefore, that attempts to measure or simulate intracellular pathway amplification have given inconclusive and sometimes opposite results (Dumartin et al., 2000; Kim et al., 2004; Marcott et al., 2014; Watts & Neve, 1996; Watts et al., 1998; Yapo et al., 2017). Therefore, it is currently unclear if the imbalance of dopaminergic binding affinity really translates into an imbalance in response to dopamine in D1 versus D2 receptor – and if it does, which receptor actually shows a stronger response.

In this study we take this problem from a top-down, functional approach instead and ask: assuming the function of the BG is to perform action selection, what is the optimal relationship between D1 and D2 sensitivity? By sensitivity, we encompass binding affinity and intracellular pathway amplification together and refer to it as “functional weight”. To address this question, we added the different D1 and D2 functional weight to the GPR model, resulting in an augmented “differential-weight” model. We then varied the value of both D1 and D2 weights independently and quantified the augmented model’s performance for soft and hard selection given each combination of weights.

## 2. Model construction

### 2.1. Input to the model

The GPR model focuses on the computations intrinsic to the BG that allow action selection. Consequently, cortical and thalamic structures are not represented in this model. Rather, it is assumed that the information entering the BG has already been pre-processed and compiled into a singledimension scalar termed “salience”. Here, each channel receives a specific, user-defined salience input that can vary over time. Additionally, since no noise is introduced in the model, the model and all its variants are fully deterministic. Consequently, for a given input and given parameters, the final outcome of a simulation should always remain the same.

**Figure 2.**
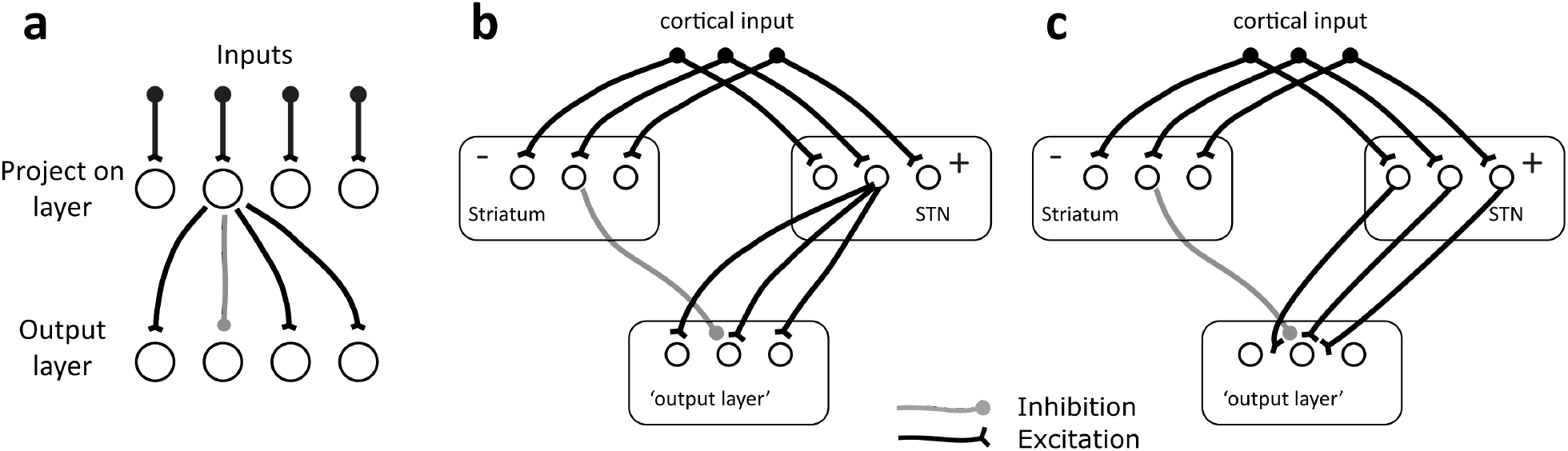
An overview of the functional architecture of the BG. **(a)** Functional architecture of a 4-channels feed-forward net implementing an off-center, on-surround computation. Only the output from the second neuron of the projection layer is explicitly shown. **(b)** Implementation of a 3-channels feed-forward net in the BG. Note that the focused inhibitory and diffuse excitatory component of the projection layer are split in two distinct neural population, respectively the striatum and the STN. **(c)** illustration of the problem of capacity scaling intrinsic to feed-forward nets. Excitatory and inhibitory projections are represented by *solid* and *grey* lines, respectively. *Figure modified from Gurney et al. (2001a)*

### 2.2. Implementation of the differential affinity of D1 and D2 receptors

In the original model, the *i*th channel receives the same salience input, *c_i_* at both D1- and D2-striatal modules. This input is then modulated by the dopaminergic input *λ* and multiplied by a shared cortico-striatal weight *w_s_* according to the following equations:

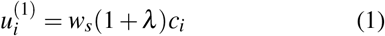

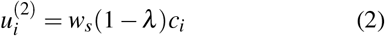

where *u_i_* is the net, ‘post-synaptic’, input to the striatum. Note that the dopaminergic modulation given by *λ* facilitates input for the striatum-D1 and attenuates it for the striatum-D2, in line with biological evidence (Surmeier et al., 2007). Since the model alteration in this study is at the level of the dopaminergic input *λ*, we focus on equations 1-2. The original model is further detailed in its entirety in the Appendix but, briefly, each nucleus in the model is designed as a set of neural units (one unit for each channel) standing for neural populations. These receive a sum of weighted inputs which is then passed through a saturating non-linear output function (equation 12 in the Appendix) to bound the output range. The connections between channels and between nuclei are as outlined in figure 1a and 2b, c.

To introduce the differential sensitivity to dopamine at D1 and D2 synapses, we introduce additional modulation weighting factors *w*_*D*1_, *w*_*D*2_, which combine with *λ* to produce receptor-type dependent modulation. Thus the equations in 1 and 2 become

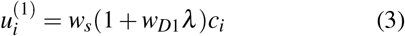

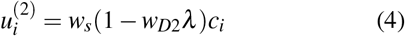

Our experiments seek to discover the effects of these modulating factors and thereby, dopaminergic affinity, on model selection. However, it is first necessary to define more precisely what we mean by ‘selection’ so that we have a quantitative measure of this function which can be evaluated for different values of *w*_*D*1_, *w*_*D*2_.

## 3. Quantifying selection capability

### 3.1. Defining selection

Recall that the BG selects via disinhibition. Thus, selection occurs on channel *i* if there is sufficient reduction in GPi output, *Y_i_*, compared to its tonic value *Y*^0^. The tonic value *Y*^0^ is the same for all channels (channel-independent) and is the output of the GPi in the absence of any input (figure 3). To quantify this reduction from the tonic value, we define a selection threshold *θ*, where *Y*^0^ > *θ* ≥ 0, and deem that selection has occurred if *Y_i_* ≤ θ. Specifically, in this instantiation of the model, *Y*^0^ = 0.17 for the parameters values specified in table 2, and we used *θ* = 0.

Now consider the case where two channels have non-zero input, so there are effectively two competing channels. To evaluate the outcome of this scenario, we run the following experimental protocol (figure 3). At *t* = 0, no input is presented to the model for a time sufficient to let tonic outputs to be expressed, and reach equilibrium. Then, at *t* = 1, one channel receives an input of salience *c*_1_. At *t* = 2, the first channel has already reached equilibrium, and a second channel receives an input *c*_2_. Because of inter-channel interactions, this second channel changes the activity of the first channel despite *c*_1_ remaining the same. The simulation finishes at *t* = 3, when both the first and second channel have reached equilibrium^1^. At this point there are three possible outcomes.

#### (1) No selection

Neither channel output has reached the selection threshold – no action is selected.

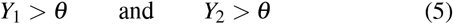

#### (2) Single channel selection

Only one channel is selected.

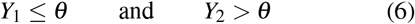

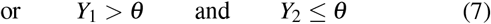

#### (3) Dual channel selection

Both channels are selected.

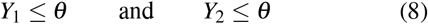

To discover the properties of selection more generally for a given set of model parameters, we repeat this experiment for all combinations of *c*_1_ and *c*_2_ between 0 and 1 with a step of 0.1. The outcome of this set of 121 simulations is reported on a 2D grid, with axes for *c*_1_ and *c*_2_, and a symbol at each grid point denoting which of the three possible outcomes occurred (as defined above); see figure 4. Panels a and b therein, show such results for typical low and high dopamine levels in the original GPR model.

**Figure 3.**
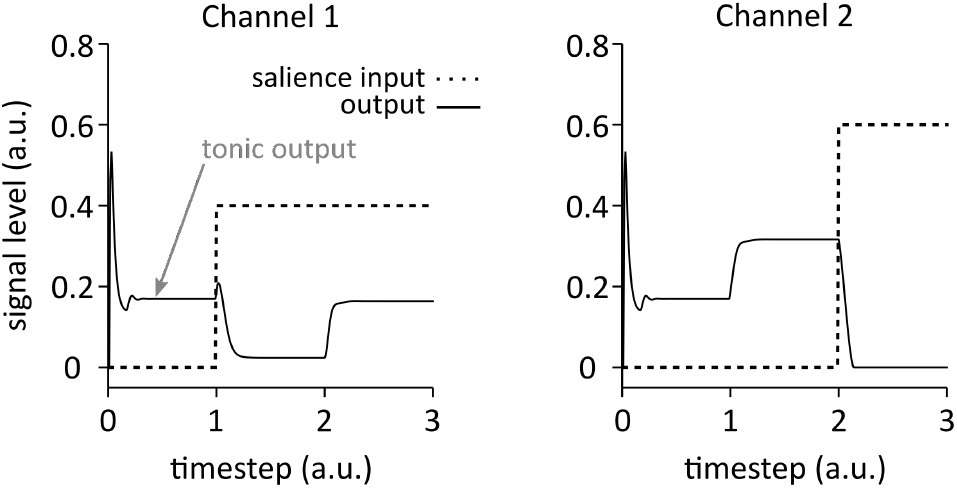
Salience input to the model and dynamics of its output. Data for channel 1 and 2 are represented in the left and right graph, respectively. Note that the change in input in channel 1 at *t* = 1 leads to a change in output value for channel 2, and *vice* – *versa* at *t* = 2. The tonic output (*i.e.* for no input at all in any channel) can be seen from *t =* 0 to *t =* 1.

### 3.2. The effects of dopamine - hard and soft selection

In order to measure selection performance of the model, it is necessary to reduce the grid of 121 experimental outcomes obtained in the previous section to a much simpler set numeric quantities and, in turn, characterise how these quantities change with dopamine modulation.

We follow here the method given in (Gurney et al., 2004) which depends on the notions of ‘hard’ and ‘soft’ selection introduced in section 1.4. The first step is to quantify precisely what these concepts mean. To do this, consider again the results for the original GPR model with low dopamine in figure 4a. Most of the experiments produced either no-selection, or single channel selection; only a handful produced dual channel selection when competition is strong at very high salience. This outcome would therefore be deemed to be more like hard than soft selection. Thus, to characterise the archetypal variant of hard selection we take this outcome to a limiting case shown in figure 4c. Here there are no instances of dual selection and the number of no-selection case is minimised (there have to be some, as disinhibition beyond threshold cannot occur for very small inputs). We refer to the outcome in figure 4a as the *hard selection template*.

Now consider the results for the original GPR model with high dopamine in figure 4b. Most of experiments produced dual channel selection which might be typical of a soft-selection regime. This is idealised in the *soft selection template* in figure 4d.

**Figure 4.**
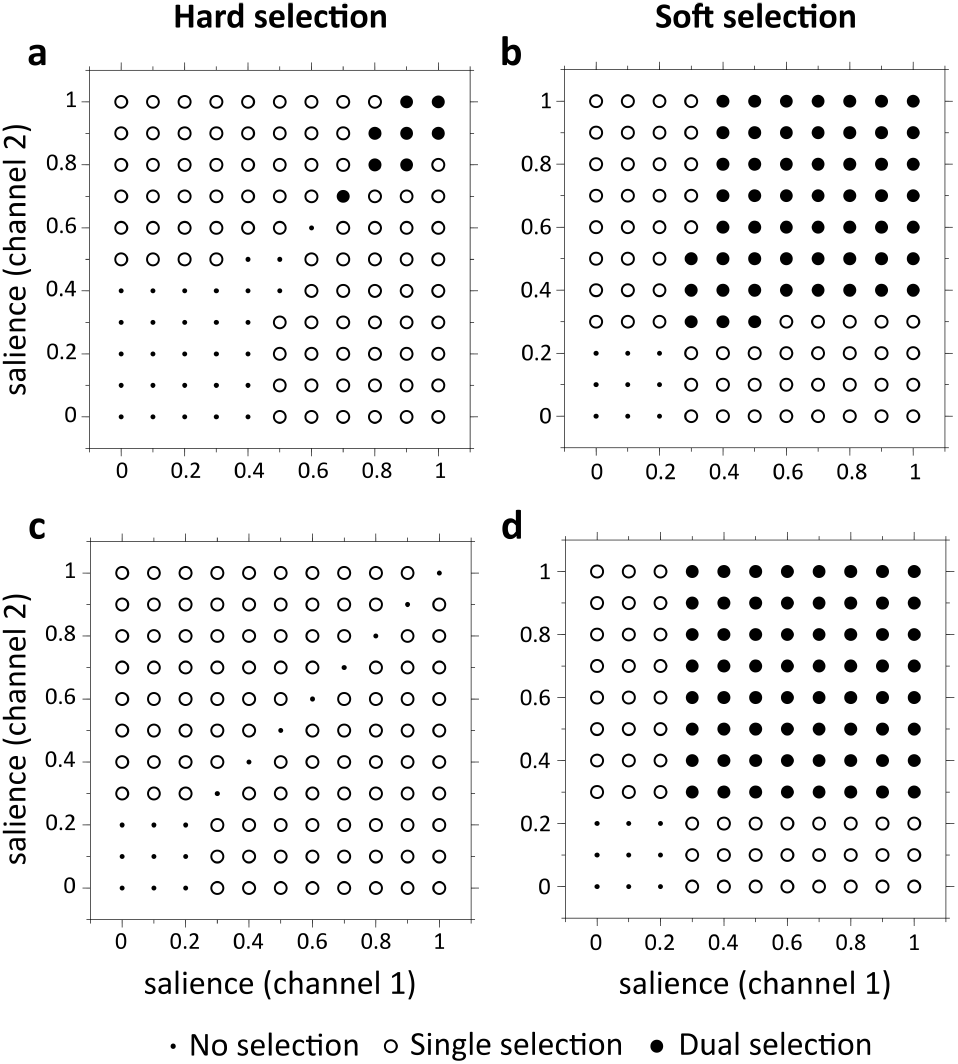
Templates for hard (left panels) and soft (right panels) selection. **(a-b)** Example of simulations outcomes from the original GPR model for hard and soft selection, using two competing channels and varying the salience input each channel receives. Hard and soft selection were performed with low (*λ* = 0.294) and high (*λ* = 0.818) dopamine levels, respectively. **(c-d)** Optimal outcomes for hard and soft action selection.

We can now use these templates to characterise the outcome of a complete set of 121 experiments by its degree of similarity with each of the templates. Thus, for each experimentally determined grid, let *N_h_* be the number of grid points with the same outcome (no-, single- or dual-selection) as the hard selection template, and put *P_h_* = 100 × *N_h_*/121. In a similar way we define *P_s_* as the degree of similarity with the soft selection template.

The values *P_h_, P_s_* will depend on the level of dopamine (as exemplified in figure 4). Thus, low levels of dopamine are likely to fit the ideal hard selection template, while higher levels are associated with soft selection. We consider this transition – from hard to soft selection – to be a feature of dopamine control of basal ganglia function (Blanco & Sloutsky, 2020; Bogacz, 2020; Costa, 2007; Costa et al., 2006; Gizer et al., 2009; Wickens et al., 2007). If the basal ganglia supports such a mechanism, then values of the parameters *w*_*D*1_, *w*_*D*2_ which will optimise this transitional control will reflect the relative biological dopamine sensitivity these parameters aim to capture. To proceed, we now require a quantitative measure of the transitional control from hard to soft selection as it depends on dopamine.

### 3.3. Quantifying dopamine-dependent selection control

We first obtain the similarity measures *P_h_, P_s_* for each level of dopamine. Dopamine is nominally parametrised by *λ* in the model, but a quantity which is perhaps easier to relate to cortico-striatal modulation is the ratio, *R_w_*, of synaptic facilitation/attenuation in the original GPR model

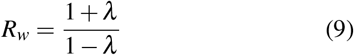

Note that *R_w_* is bounded below by 1 since *λ* is bounded below by 0, but it has no theoretical upper bound. Then, for a given dopaminergic ratio, *R_w_*, we run a grid set of experiments and obtain *P_h_* (*R_w_*) and *P_s_* (*R_w_*). We can then plot these functions as shown in figure 5a.

**Figure 5.**
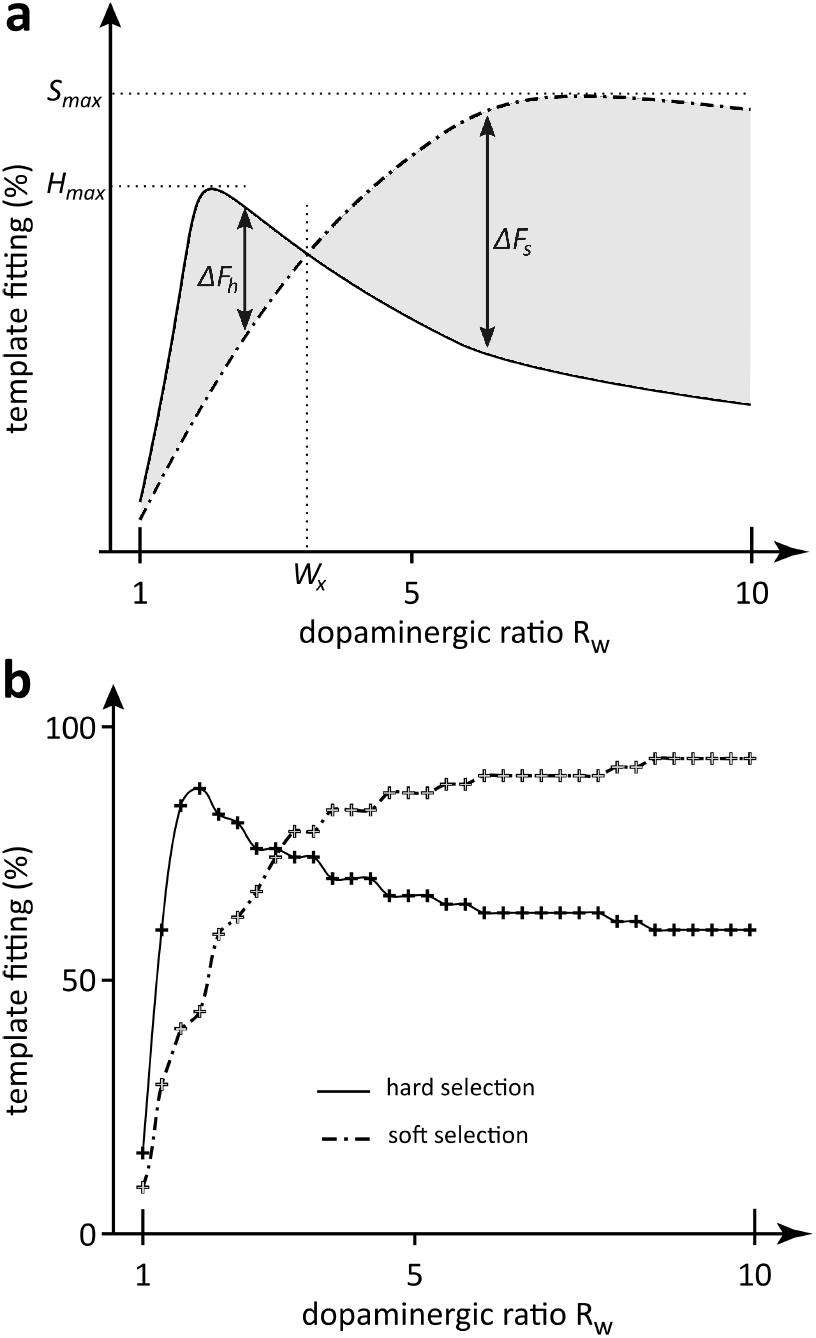
Template fitting as a function of *R_w_*. **(a)** Cartoon representing the variables used to define the template fitting functions’ main features. Hard and soft selection templates *P_h_* and *Ps* are represented in *solid* and *dashed* lines, respectively. **(b)** Example of template fitting function for the set of outcome of the original GPR model. Only a third of the data points are represented here for each functions, for visibility. *Panel a modified from Gurney et al. (2004).*

In an ideal selection machine which allows control of the hard/soft selection via the parameter *R_w_*, we might expect the following characteristics. First there will be well defined regimes for hard and soft selection, separated by some crossover point at *R_w_* = *w_x_*. For 1 ≤ *R_w_* < *w_x_*, hard selection will dominate and so *P_h_* (*R_w_*) > *P_s_* (*R_w_*); for *w_x_* ≤ *R_w_* soft selection prevails and so *P_h_* (*R_w_*) < *P_s_* (*R_w_*). This is shown in figure 5a. Notice that the observed behaviour of the GPR model (figure 5b) complies with this criterion. The degree to which the two regimes are well defined and contrasted may be captured in any number of ways, but we adopt a feature-based approach as shown in the figure. The features are: the maximum value of *P_h_* in the hard selection regime, *H_max_*; the corresponding maximum for soft selection, *S_max_*; the mean valueΔFh of the difference *P_h_* – *P_s_* for hard selection and, the mean value Δ*F_s_* of *P_s_* – *P_h_* up to some practical maximum value of *R_w_* = 10. While this maximum value is somewhat arbitrary, we expect that physiological constraints will also limit any biological correlate of this ratio.

In an ideal selection machine we would expect all these features to have as large a value as possible. In addition we might expect the transition point *w_x_* to be significantly larger than 1, otherwise there is little room for adopting the hard selection regime under noisy control of the parameter *R_w_*. We therefore consider *w_x_* as a member of the feature set describing selection.

In order to facilitate comparison with the original GPR model it is useful to invoke the ratios of the feature values for any new model with respect to those for the GPR model. Thus if *f* is one of the five features {*w_x_*, Δ*F_h_*, Δ*F_s_*, *H_max_*, *S_max_*} put *r* = *f/f^G^* where *f^G^* is the GPR model value. We also log-transform defining 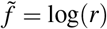; for example 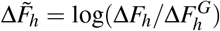, where 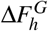 is the value for the GPR model. Then 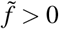 implies a feature value larger than that of the GPR model, which, by dint of the way the features were defined, means a better hard/soft selection performance

The features can then be used to define a metric, *Q* which measures the quality of the transitional control between the two regimes. Thus, let 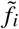 be the (log-transformed) ith feature (1 ≤ *i* ≤ 5. Then assuming all features are well defined, and that no *r_i_*, = 0, put 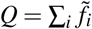. The GPR model therefore has *Q* = 0 and so positive/negative values of *Q* imply a performance better/worse than the baseline of GPR. If, however, any model feature is not defined or zero, then *Q* is also not defined. This indicates a failure of the model to display sensible hard/soft selection behaviour. Full details of the features and their definition can be found in Gurney et al. (2004)

## 4. Results

A series of experiments were done by sampling *w*_*D*1_, *w*_*D*2_ on a grid over their respective ranges of values. Note that we must have *w*_*D*2_ *λ* ≤ 1 since values larger than this can allow for and overall change the sign of the cortico-striatal input in equation 4, which is biologically implausible. Since we ran our simulations with *R_w_* ≤ 10, rearranging equation 9 gives us *λ* ≤ 9/11 and so we must have *w*_*D*2_ ≤ 11/9. *w*_*D*1_ is not bounded as such, but performance peaked at values well under *w*_*D*1_ = 10 which therefore was a practical upper bound for our simulations. Thus, we had 0 ≤ *w*_*D*1_ ≤ 10 and 0 ≤ *w*_*D*2_ ≤ 11/9. We ran the experiments with 401 and 50 sampling points for parameter *w*_*D*1_ and *w*_*D*2_, respectively, to ensure at least a sampling point every 0.0250 step. This resulted in 20,050 complete evaluations of the model, each yielding a set of features and *Q* value indicating its performance relative to the original GPR model.

### 4.1. The sensitivity weightings can radically change performance

Some representative template fitting functions are given in figure 6 with results for the original GPR model shown in panel a. Panel b shows an instance where there is no hard-soft transition within the domain explored and so the *Q* metric is undefined. In contrast, panels c and d show two cases of successful models with *Q* > 0 and *Q* < 0, indicating better and worse performance than the original model, respectively. The detailed characteristics and feature set values of each model illustrated in figure 6 is available in table 1.

**Figure 6.**
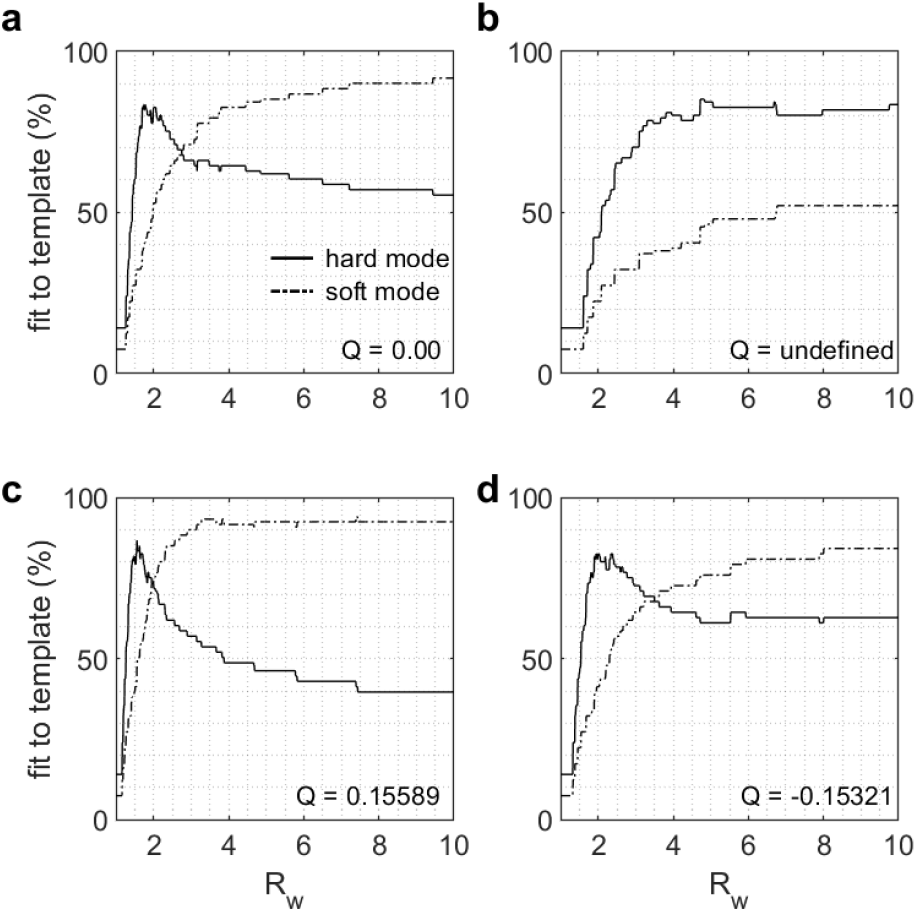
Sample of representative template fitting functions for different versions of the augmented model. The weights of each version evaluate at **(a)** *w*_*D*1_ = 1, *w*_*D*2_ = 1 (the original model), **(b)** *w*_*D*1_ = 0.6, *w*_*D*2_ = 0.4, **(c)** *w*_*D*1_ = 1.5, *w*_*D*2_ = 0.6, **(d)** *w*_*D*1_ = 2, *w*_*D*2_ = 1.2. The solid and dashed lines represent *P_h_* and *P_s_,* respectively.

**Table 1.**
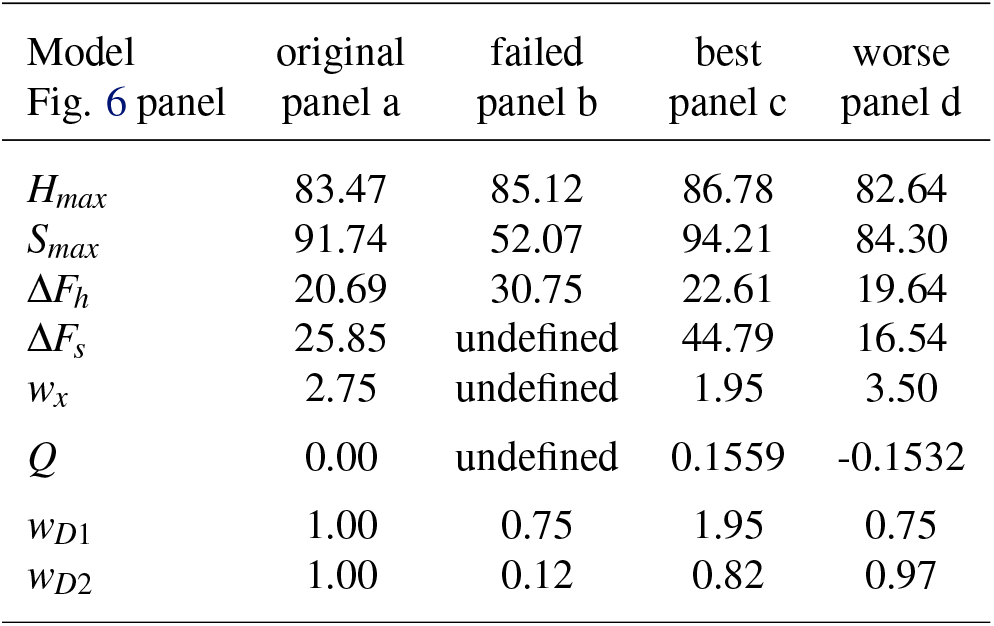
Feature set, merit *Q* and parameter values of each model from figure 5.

The better performance of the model in panel c is due to the marginally greater fit to the optimal hard and soft templates displayed in figure 4c-d (*H_max_*, *S_max_* in table 1). The transition between hard and soft selection modes is also steeper in panel c, indicating a better Δ*F_h_* and the difference between hard and soft selection curves is also wider on average after the transition point (Δ*F_s_*). However, the transition point *w_x_* is closer to 1 for the model in panel c than in the original GPR model, illustrating that a positive *Q* value does not necessarily mean that all the underlying features behave better than the original model.

Panel d illustrate how a model that is successful, but less efficient than the control model, may behave. In this particular case, even though *w_x_* is further away from 1 (table 1), hard and soft selection regimes (*H_max_* and *S_max_*) are much less efficient and the (normalised) difference between the hard and soft curve is smaller before the transition (Δ*F_h_*) as well as after the transition (Δ*F_s_*).

Overall, these examples illustrate the wide variety of behaviours that the augmented model can exhibit as a function of the *w*_*D*1_, *w*_*D*2_ parameter value used. To obtain a more comprehensive view of the effect these parameters have on the augmented model, we then assessed how each of the features vary as a function of each *w*_*D*1_, *w*_*D*2_ pairs.

### 4.2. Increased sensitivity in D1-striatum gives robust enhanced performance

Figure 7 shows each of the features, as well as the augmented model’s merit *Q* as we vary *w*_*D*1_ and *w*_*D*2_. Note that, here, the features are expressed in as a function of the original model (*i.e.* 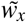 instead of *w_x_*) to allow for better comparison, as discussed in section 3.3. Therefore, a feature value above/below 0 indicates better/worse performance than the original GPR model for the corresponding *w*_*D*1_, *w*_*D*2_ pair.

**Figure 7.**
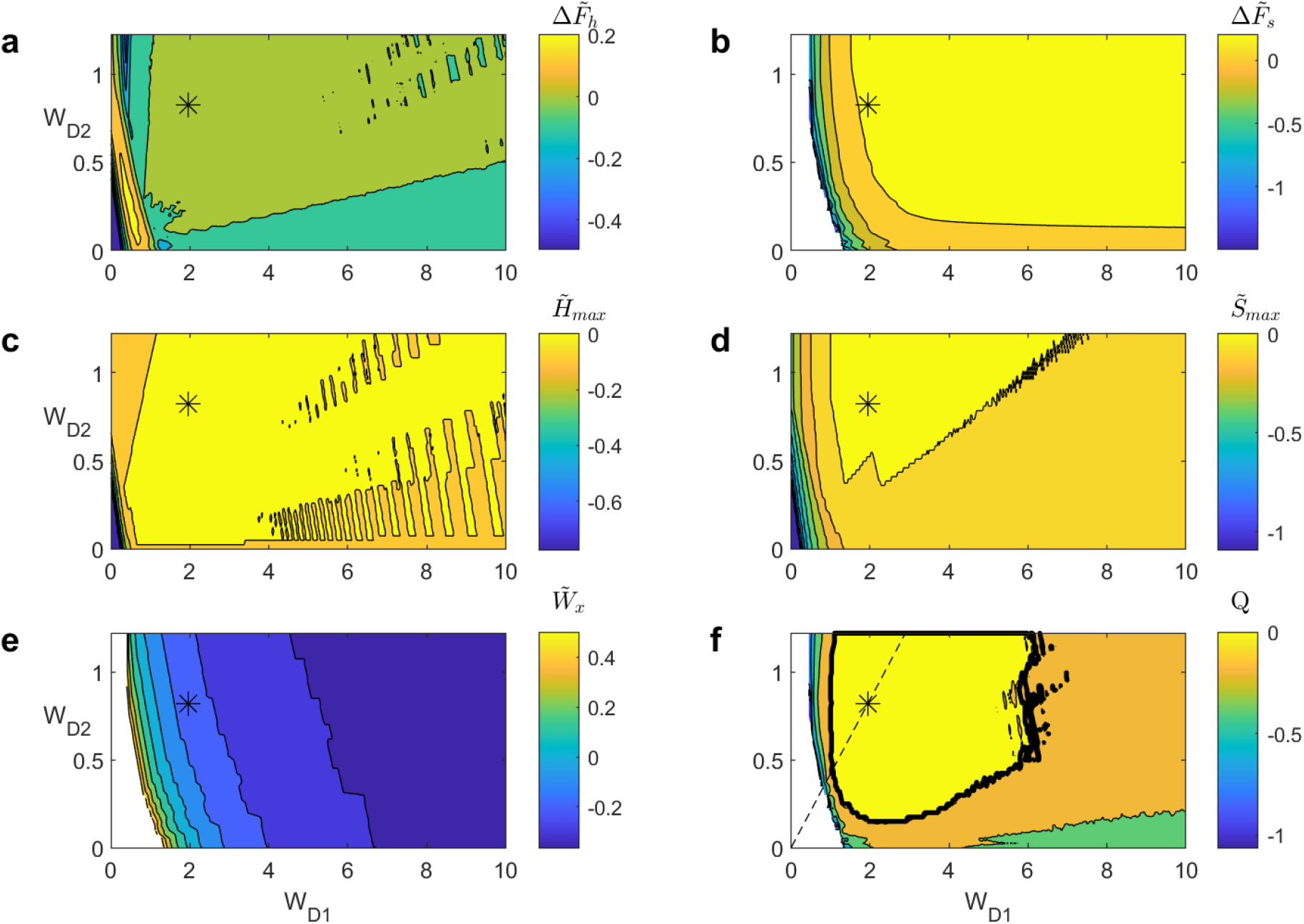
Evolution of **(a)** 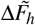, **(b)** 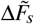, **(c)** 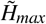, **(d)** 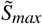, **(e)** 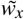 and **(f)** *Q* as a function of *w*_*D*1_ and *w*_*D*2_ for the different versions of the augmented model. Cold and warm colors represent low and high values, respectively. Empty (*i.e.* white) values mean that the variable is undefined. The * mark on each panel indicates the location of the best *Q* value. The solid black line in **(f)** indicates the regions for which the augmented model is better than the control model (Q > 0) and the dashed line indicate the optimal ratio *w*_*D*1_ /*w*_*D*2_ = 2.369 obtained from the ratio analysis (see section 4.3).

A first observation is that an overly small *w*_*D*1_ or *w*_*D*2_ value leads to a failed model, because 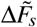 and 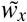 are note well defined (figure 7b, e, f). This is because small *w*_*D*1_ and *w*_*D*2_ values effectively shrink the effect of dopamine on the cortico-striatal input, eventually forcing the model to behave as it does when *λ* is always close to 0 (see equations 3-4). This hinders the model’s ability to switch to soft selection at high *λ* values, leading to models behaving similarly to what we observe in figure 6b.

The opposite occurs as *w*_*D*1_ and *w*_*D*2_ take very large values. The *λ* modulation on cortico-striatal input is magnified, forcing the model to quickly transition to a soft selection mode. This can be seen by looking at 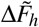 in figure 7a, which reaches a (positive) peak value for *w*_*D*1_ < 1.5 to then drop below 0 as *w*_*D*1_ keeps increasing. Similarly, we see that 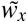 initially takes high, positive values, and then drops to negative values as *w*_*D*1_ increases as well, indicating that the transition occurs for lower *R_w_* values than in the original model (figure 7e).

The remaining three features, 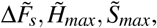 display roughly identical patterns (figure 7b-d). Excluding the undefined region for 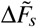, performance first increases rapidly to then quickly saturate at values ≥ 0 as *w*_*D*1_, *w*_*D*2_ increase as well. This behaviour for 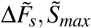 again outlines that large *w*_*D*1_, *w*_*D*2_ values facilitate soft selection by magnifying the effect of *λ* on cortico-striatal input.

Taking into account the behaviour of each feature, it becomes clear that a model that behaves best is one that will take intermediate *w*_*D*1_, *w*_*D*2_ values. This enables striking a good balance between a sharp and clean hard-soft selection transition, while allowing the model to reach the optimal soft selection regime only available at high-dopamine levels. This is illustrated by the behaviour of the *Q* value, which incorporates all of these features into a single metric (figure 7f). We see that *Q* is first undefined because 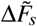 and 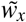 are undefined as well. It then quickly increase to a reach a positive region around *w*_*D*1_ = 1 and *w*_*D*2_ = 0.2. It then takes its maximal value (*Q* = 0.15589) at *w*_*D*1_ = 1.95 and *w*_*D*2_ = 0.75 (table 1, figure 6c), indicated by a * in figure 7a-f. Finally, its value decreases back to negative values as *w*_*D*1_, *w*_*D*2_ keep increasing, outlining the detrimental effect these larger weights have on 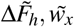.

Overall, these results outline that the augmented model can perform better than the original model given appropriate *w*_*D*1_, *w*_*D*2_ values. Further, there is a fairly extensive range of these weightings for which this is true (see the region for which *Q* > 0 in figure 7e). Unsurprisingly, overly small *w*_*D*1_, *w*_*D*2_ values can lead to a failure of the model because it prevents any soft selection from occurring. However, extremely large values do not lead to model failure, although they are clearly sub-optimal and thus undesirable. Interestingly, all values of *Q* > 0 occur for *w*_*D*1_ > 1; in contrast *w*_*D*2_ < 1 can result in *Q* > 0. In fact, the *w*_*D*1_, *w*_*D*2_ pair that led to the best performance have *w*_*D*1_ > 1, *w*_*D*2_ < 1, suggesting that the selection pathway benefits from cortico-striatal input facilitation while the control pathway benefits from its attenuation.

### 4.3. There is a weak dependence of *Q* on the D2 to D2 weight ratio

Since *w*_*D*1_ and *w*_*D*2_ are applied to the same input values in each population of the striatum (see equations 3-4), it may be that their ratio, rather than their absolute value matters most for the purpose of efficient computation. Therefore, we assessed the merit of the augmented model as a function of a ratio *w*_*D*1_ /*w*_*D*2_.

We observe that a ratio under 1 results in a sub-optimal merit for the augmented model compared to the original model (*i.e. Q* < 0; figure 8). From thereon, marked improvements from the original GPR model can be observed throughout ratios ranging from 1 to 5 (*Q* > 0), and the best *Q* value we observed was for *w*_*D*1_/*w*_*D*2_ = 2.369, for which we had *Q* = 0.15589. Beyond around 5, Q decreases as the weight ratio increases. Thus, *Q* ‘plateaus’ in the range 1-5, suggesting that there is no particular ratio value for which the augmented model displays a superior performance. This is in line with the results of the previous section, where there was an extensive region of improved performance.

**Figure 8.**
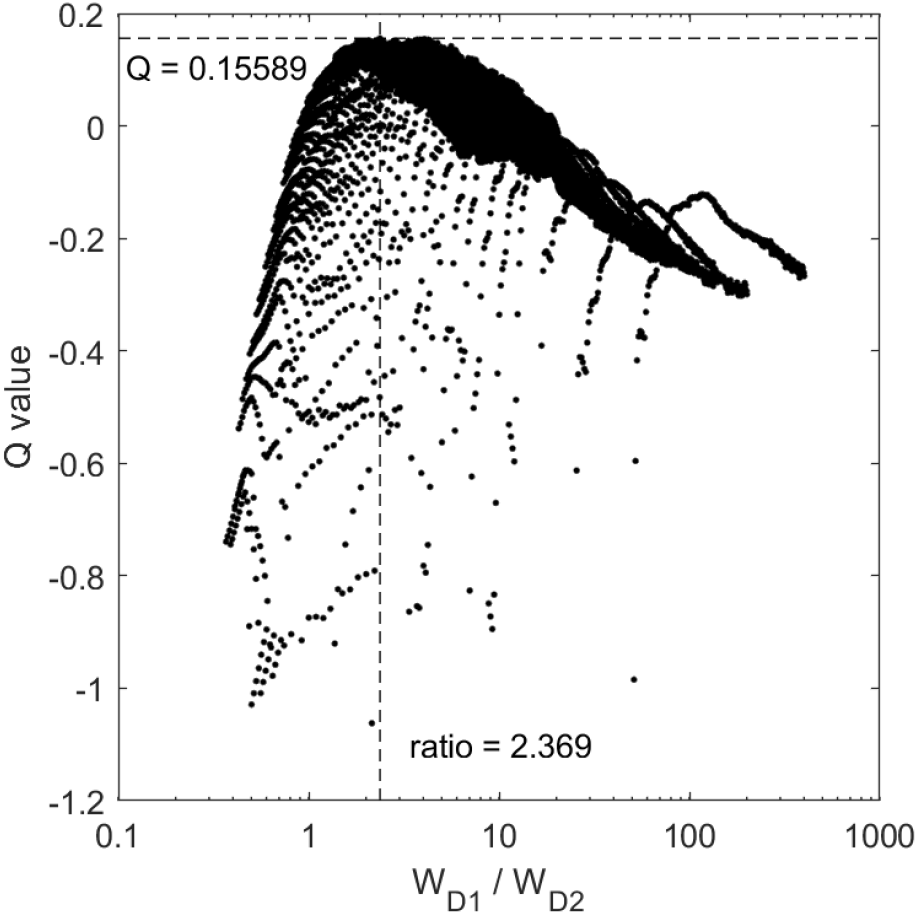
Model merit *Q* as a function of *w*_*D*1_ /*w*_*D*2_ ratio. The x-axis is in a logarithm base 10 scale. The dashed lines indicate the coordinates for the best *Q*

However, not all *Q* values in the “optimal” range of ratios (1 to 5) are positive, demonstrating that the augmented models’ merit depends not only on the *w*_*D*1_ /*w*_*D*2_ ratio value, but also particular values of each weight. This is likely because the model behaves more poorly as *w*_*D*1_ and *w*_*D*2_ get closer to 0, even if it lies on the best-ratio line (see dashed line in figure 7f).

## 5. Discussion

From the behaviour of model performance as we varied *w*_*D*1_, *w*_*D*2_, we observe that both large and small weight values present advantages. Small weight values allow the model to express a clean hard selection mode as well as a sharp transition from hard to soft selection modes (figure 7a). In contrast, large weights values enable the model to reach for an optimal soft selection mode that would otherwise require a greater dopaminergic modulation (figure 7b,d). Consequently, the optimal model we observe has intermediate weight values, striking a balance between those two modes of behaviour. Critically, the optimal model performs better than the original GPR model and has non-equal weights *w*_*D*1_, *w*_*D*2_, thus indicating that differential sensitivity to dopamine is a benefit to selection. In addition, the model behaviour is robust to small changes in these weights. Finally, we observe that this improved performance is only weakly related to the relative *w*_*D*1_ /*w*_*D*2_ ratio, as it also depended the absolute value these weights take. Particularly, the model benefits from *w*_*D*1_ > 1. There was also some evidence that having *w*_*D*2_ < *w*_*D*1_ benefited the model, as very few augmented models with a weight ratio below 1 showed improved performance from the original model.

How do the optimal weights *wD*1 and *wD*2 observed here compare to physiological observations? Two approaches can be used in order to experimentally determinate the functional connectivity between neurons. First, one can record the electrophysiological response of the post-synaptic neuron following artificial stimulation of the pre-synaptic neuron (Chuhma et al., 2011). However, such approach is technically challenging, and the dopamine effect on striatal neurons depends on the overall state of the network (Surmeier et al., 2007). For instance, striatal neurons may be up-state or down-state or the drive of cortical input may vary. Therefore, there is to our knowledge no study that directly assesses the functional connectivity of dopaminergic neurons in the striatum.

The second approach consists in quantitatively determining the functional contribution of each of the steps involved in synaptic transmission that contribute to the complete post-synaptic response. There are three such contributions: the proportion of D1 and D2 receptors expressed; their binding affinity; and intracellular signal transduction following receptor activation.Generally, quantifying D1 and D2 sensitivity has proved a challenging question, and the answer likely differs depending on the timescale and temporal profile of the dopaminergic signal, and state of the post-synaptic neuron (Cumming, 2011; Kenakin, 2013; Marcott et al., 2014; Richfield et al., 1989; Skinbjerg et al., 2012; Yapo et al., 2017). Here we review, in the context of the current model, some of the data pertaining to each of the three possible mechanisms underlying dopamine sensitivity,

Perhaps the least challenging contribution to estimate is that of the proportion of D1 and D2 receptors expressed. Several studies suggest that D1 receptors are about twice as common as D2 in the striatum (Dewar et al., 1997; Roseboom & Gnegy, 1989; Wanderoy et al., 1997) thereby enhancing relative sensitivity in the D1 pathway. This is consistent with our modelling results showing that greater D1 sensitivity (*w*_*D*1_ > *w*_*D*2_) benefits action selection computation.

Regarding binding affinity, previous work evaluated 20% and 90% affinity for D1 and D2, respectively (Richfield et al., 1989). However, these estimates can suffer from methodological limitations: D1 and D2 both show high and low affinity states depending on their intracellular G-protein state, *in vivo* and membrane preparation experiment gave occasionally inconsistent results, or the binding affinity of D2 varies based on factors such as amphetamine regulation (see Cumming (2011); Skinbjerg et al. (2012) for reviews). Therefore, while the reported results on affinity appear to be at odds with our result, their full interpretation remains unclear.

Concerning intracellular signal transduction, both D1 and D2 pathways lead to a phosphorylation of DARPP-32, a molecule that is sometimes considered as the final effector of D1 and D2 intracellular pathways (Lindskog et al., 2006; Surmeier et al., 2007; Svenningsson et al., 2004). Phosphorylation of DARPP-32 is increased following D1 receptor binding, but decreased following D2 receptors binding (Nishi et al., 1997; 2000), which is in line with the functionally opposite effect of D1 and D2. Critically, an increase of 6-folds in DARPP-32 phosphorylation can be observed for D1 activation, while a 2-folds decrease is observed following D2 activation (Nishi et al., 1997; 2000). This would suggest a sensitivity contrast in line with our results. However, measurements of intracellular signaling have sometimes provided conflicting accounts, adding to the uncertainty on the D1 and D2 functional weights (Dumartin et al., 2000; Kim et al., 2004; Marcott et al., 2014; Watts & Neve, 1996; Watts et al., 1998; Yapo et al., 2017).

Taken together, the larger proportion of D1 receptors expressed and the greater intracellular amplification of their effect could facilitate a greater D1-sensitivity, despite some evidence for a D2 binding affinity advantage. Thus, our prediction, based on the action selection hypothesis extended to include dopaminergic control of selection regime, is that there is an overall increased sensitivity to dopamine in the selection (D1) pathway, compared to that in the control (D2) pathway.

There are several aspects of the BG physiology that may impact cortico-striatal input modulation and that were not modelled here. For instance, the up- and down-state condition of striatal neurons alters the effect of dopamine on its post-synaptic targets (Arbuthnott & Wickens, 2007). While some broad aspects of striatal neuron state are captured in the nonlinear output function (see appendix), this phenomenon is complex and would require modeling at a lower level of description.

We have also not modelled the effects of phasic dopamine release which is believed to play an important role on learning (Schultz, 1998). Rather, we have focused on tonic dopamine levels that are believed to be more important for selection functionality (Groves et al., 1994).

Dopamine sensitivity at the level of striatum is physiologically grounded in many complex and interacting mechanisms whose overall effect is intractable just now. We have used a high level computational model, and an extension of the action selection hypothesis (to include dopaminergic control of hard/soft selection), to predict the relative sensitivity to dopamine at D1- and D2-dominant sites in striatum. We have shown that selection can benefit from differential rather than identical, sensitivity therein, and that this benefit is seen with D1-favourable emphasis in such sensitivity.

## A. Appendix

The parameter values employed are listed in table 2. All channels receive a scalar input *u* defining an activity function *a* over time using an ordinary differential equation

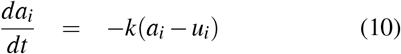

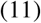

where *i* indicates the ith channel, *k* is the decay constant, *t* is time. Since at equilibrium *da_i_/dt* = 0, the equation converges to *a_i_* = *u_i_*. *a_i_* is then passed through a piece-wise output function *y_i_* to avoid runaway activity

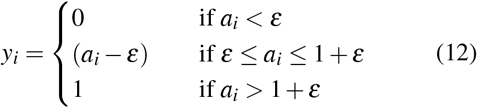

where ε is a constant threshold whose value depends on the nucleus, as defined below.

### Striatum

Although the mathematical implementation of the striatum is already detailed earlier, it is repeated here for convenience. In the original GPR model, the striatum D1 and striatum D2 receive the same salience input *c_i_* from the cortex, which is then modulated by the dopaminergic input *λ* and weighted by a shared striatal weight *w_s_*

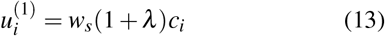

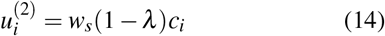

The output of the striatum D1 and striatum D2 is denoted 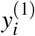 and 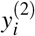, respectively, after applying equations 10 and 12 - with a shared threshold ε^−^ for the output function (see equation 12). Note that for *λ* = 0 (no dopamine), the dopaminergic modulation (1 + *λ*) and (1 – *λ*) reduce to 1. Finally, eq. 13-14 are altered in the augmented model, as indicated in eq. 3-4.

### STN

The input of the STN includes excitatory projections from the cortex and inhibitory projections from the GPe (figure 1). Let us denote the output of GPe as *Y_i_* and its weight when contacting STN (GPe→STN) be *w_g_*. Let us denote the cortex→STN weight as *w_t_*. The input of the STN is then defined as

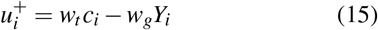

The output of the STN is then denoted 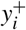 and the threshold of its output function is ε^+^.

### GPe

Let 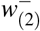 and *w*^+^ be the connection weight between the striatum D2→GPe and the STN→GPe, respectively. The GP receives diffuse excitatory projections from the STN. Consequently, the drive from this nucleus is formalized as the sum of the output of all channels such as 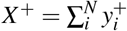 where *N* is the number of channels in the model. The input to the GPe is then defined as

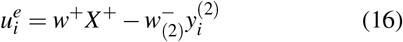

Its output is denoted *Y_i_* and the output function threshold is denoted *ε_e_*. Importantly, Y_i_ is also the final output of the model for channel *i*.

### GPi

The GPi also receives diffuse excitatory input from the STN, as well as inhibitory input from the striatum D1 and the GPe. Let us define 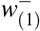 and *w_e_* the connectivity weights for the striatum D1→GPi and GPe→GPi, respectively. The input to the GPi is then defined by

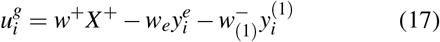

Its output is denoted 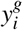 and the output function threshold is ε_*g*_.

### Parameters value

Table 2 recapitulates all the parameters that are employed here as well as the value they take, which was identical for the original and the augmented models. A rationale of most parameters of the original model can be found in Gurney et al. (2001a) and Gurney et al. (2001b).

The main criterion used to define the model’s connectivity weights was to tune the activity variables according to the output functions. In order to achieve that, *w_s_*, *w_t_*, *w_g_,* 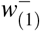 and 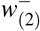 were set to 1. For theoretical reasons, *w*^+^ had to be less than 1, and was therefore set at 0.9 (see Gurney et al. (2001b)). These values were used for both the original and augmented models. Finally, because the original model behaved poorly for high values of *w_e_,* this parameter was evaluated at 0.3 (Gurney et al., 2001b). The decay constant *k* was evaluated at 25.

All the models, including the original one, contain *N* = 6 channels. However, in the simulation, only 2 of them receive salience input, that is, we always have *c*_3–6_ = 0. This allows keeping the set of potential outcomes simple while still exposing the effect of channel-to-channel competition. The role of the remaining 4 channels is to model the tonic inhibition from quiescent channels via the diffuse projections of the STN. Previous work on the original GRP model also contain 4 quiescent channels for the same reasons.

**Table 2.**
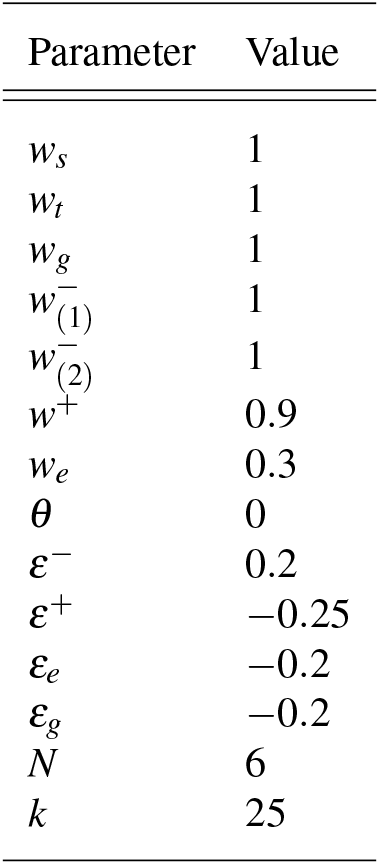
Parameters value for the GPR model.

The parameter *R_w_* will range from 1 to 10, because when *λ* = 0 we have *R_w_* = 1 and spanning this parameter until 10 allows varying dopamine levels enough for the model to fully and efficiently express both hard and soft selection (Gurney et al., 2004). Moreover, since the dopaminergic modulation of the striatum D2 is formalized as (1 – *λ*), we want to avoid *λ* > 1 because it would shut down the control pathways. Since we have *λ* = (*R_w_* – 1)/(*R_w_* + 1), *R_w_* = 10 gives us *λ* = 9/11 at maximum. This is important to bear in mind, because it implies that one of the new parameter added to the augmented model, *w*_*D*2_, cannot evaluate at 11/9 or more, because when dopamine reaches the maximum level during simulations, the control pathway would be shut down, or even inverted, which is biologically implausible.

## Footnotes

1 Since the outcome only depends on the state of the system at *t* = 3, there is no specific reason to start *c*_1_ and *c*_2_ at different times, except to illustrate between-channel interactions

